# Signal, bias, and the role of transcriptome assembly quality in phylogenomic inference

**DOI:** 10.1101/2020.07.23.216606

**Authors:** Jennifer L Spillane, Troy M LaPolice, Matthew D MacManes, David C Plachetzki

**Affiliations:** University of New Hampshire, Molecular, Cellular, and Biomedical Sciences Department, Durham, NH 03824, USA; Hubbard Center for Genome Studies, University of New Hampshire, Durham, NH 03824, USA

## Abstract

The empirical details of whole transcriptome sequencing and assembly have been thoroughly evaluated, but few studies have addressed how user-defined aspects of the assembly process may influence performance in phylogenomic analyses. Errors in transcriptome assembly could affect ortholog prediction, alignment quality, and phylogenetic signal. Here we investigate the impacts of transcriptome assembly quality in phylogenomic studies by constructing phylogenomic data matrices from alternative transcriptome assemblies representing high-quality and intentionally low-quality assembly outcomes. We leveraged a well-resolved topology for craniates to apply a topological constraint to our analyses, providing a way to quantify phylogenetic signal. Craniates are amply represented in publicly available raw RNA-seq repositories, allowing us to control for transcriptome tissue type as well. By studying the performance of phylogenomic datasets derived from these alternative high- and low-quality inputs in a controlled experiment, we show that high-quality transcriptomes produce richer phylogenomic datasets with partitions that have lower alignment ambiguity, less compositional bias, and stronger phylogenetic signal than low-quality transcriptome assemblies. Our findings demonstrate the importance of transcriptome assembly in phylogenomic analyses and suggest that a portion of the uncertainty observed in phylogenomic studies could be alleviated at the assembly stage.

## Introduction

The genomics revolution has resulted in a transformation of the approaches that scientists use to estimate phylogeny by vastly increasing both the number of available independent genetic markers (Blair et al. 2002; Dopazo et al. 2004), as well as the number of taxa included in phylogenetic analyses (Dunn et al. 2008). However, for taxa that remain largely unrepresented in publicly available datasets, generating a large number of genetic markers, often accomplished as part of a *de novo* whole genome sequencing project, continues to be a challenge. Transcriptome sequencing is typically a less challenging method of generating genome-wide marker data (although see Vijay *et al.* 2013) in that it requires fewer sequenced reads and is therefore less expensive than whole genome sequencing. In addition, transcriptomes perform comparably to genomes in phylogenomic studies when used with robust methods of ortholog identification (Cheon et al. 2020). For these reasons, data derived from transcriptome assemblies have become widely used in phylogenomic studies and have come to represent a mainstream approach to phylogenetic reconstruction (Chen et al. 2015; Reich et al. 2015; Kutty et al. 2018).

The generation of a phylogenomic data matrix is a complex and critical process, as biases introduced at this point can propagate in downstream analyses in unpredictable ways. Phylogenomic data matrices are composed of multiple (often hundreds of) partitions, alignments of orthologous loci that have been filtered and concatenated together, which makes these data matrices highly dimensional. In addition, phylogenomic datasets are often comprised of an agglomeration of data from multiple research groups that may have leveraged different sequencing and assembly strategies. Therefore it is not surprising that there are still many questions concerning the best practices related to the generation and application of these massive new datasets to phylogenomics (Wen et al. 2015; Mckain et al. 2018; Yu et al. 2018). Many researchers have addressed questions related to the most appropriate modeling schemes for different partitions of the data matrix (Philippe et al. 2005; Blanquart and Lartillot 2008; Lanfear et al. 2014; Whelan et al. 2015; Feuda et al. 2017; Wang et al. 2018). Some have considered the impact of incomplete lineage sorting in phylogenomic reconstruction and have leveraged this property of recently diverged lineages to inform species trees (Liu et al. 2010; Zhang et al. 2018). Others have sought to examine differential phylogenetic signal among partitions in order to maximize phylogenomic performance (Borowiec et al. 2015; Simion et al. 2017). Increasingly, researchers have added the additional step of recoding the amino acid data matrix in an attempt to account for saturation and compositional heterogeneity (Masta et al. 2008; Lasek-Nesselquist 2012; Feuda et al. 2017; Marletaz et al. 2019 though see Hernandez AM, Ryan JF, personal communication, https://doi.org/10.1101/729103, last accessed July 14, 2020). While each of these issues is critical to consider in phylogenomic studies, collectively they deal with aspects of the analyses that occur after the primary phylogenomic data matrix has been assembled. In most cases, biases introduced during the generation of the matrix itself are not explicitly addressed and may persist in influencing downstream inferences.

Whole transcriptome sequencing is itself a relatively new technology, having gained widespread popularity only in the past decade (Wang et al. 2009). Confusion still exists about many aspects of RNA-seq and its analysis – from how deeply to sequence, to how to gauge the quality of the transcriptome assembly. While many genomics studies have investigated methodological impacts of read trimming (MacManes 2014; Mbandi et al. 2014), error correction (Le et al. 2013; MacManes and Eisen 2013; Song and Florea 2015), different approaches to transcriptome assembly (MacManes 2018), and quality assessment (Li and Dewey 2011; Li et al. 2014; Smith-Unna et al. 2016), researchers using transcriptome assemblies for phylogenomic applications have been slow to adopt many of the recommendations of the genomics community (but see Parks *et al.* 2017). Indeed, it is still commonplace in phylogenomics analyses to treat the phylogenomic data matrix as a black box, with little focus, or in some cases transparency, regarding the nature and quality of sequence reads generated or the assemblies used as input.

To date there has been no empirical study of how transcriptome assembly quality may affect downstream phylogenomic analyses, although many impacts are possible. Poor-quality assemblies may alter the accuracy of ortholog prediction, alignment quality, and phylogenetic signal. We predicted that in phylogenomic analyses, poor-quality assemblies would result in differences in the number and identity of orthogroups as well as differences in the quality of the partition alignments compared to those from higher-quality transcriptomes. Here we examine the effects of transcriptome assembly quality on these metrics. We use a well-characterized quantitative metric (*TransRate* score; Smith-Unna et al. 2016) to evaluate transcriptome assemblies and systematically to construct two separate phylogenomic datasets: one of high quality and one of intentionally low quality. We then perform identical phylogenetic analyses on each dataset, identifying discrepancies between them and assessing the extent to which these differences affect phylogenetic inferences. We find that high-quality transcriptomes produce larger phylogenomic datasets with partitions that have less alignment ambiguity, weaker compositional bias, and stronger phylogenetic signal than datasets from low-quality transcriptome assemblies. Our results indicate that a portion of the uncertainty in phylogenomic studies likely stems from issues related to the initial assemblies used in preparing phylogenomic data matrices.

## Results

### Datasets chosen based on TransRate scores show little variation in BUSCO score

Our study design controls for several factors that could preclude direct comparison between empirical outcomes in phylogenomic analyses. We focus on the craniate phylogeny because there is little debate about the major relationships within the group and because RNA-seq read data are available from the same tissue type (liver) for a wide range of taxa. The read sets used in this study ranged in size from 13.7 million read-pairs (*Calidris pugnax*) to 46.4 million read-pairs (*Ambystoma mexicanum*). We prepared two separate datasets from the same read sets, one of high quality and one of low quality using the Oyster River Protocol (MacManes 2018).

The high-quality dataset based on overall *TransRate* assembly scores had an average *TransRate* score of 0.47236 (ranging from 0.23542 to 0.68372), while the low-quality dataset’s average *TransRate* score was 0.15943 (ranging from 0.09216 to 0.25281). The read-pairs assembled into significantly fewer transcripts in the high-quality dataset compared to the low-quality dataset (*P* < 0.001, Figure 2A), with an average of 178,473 and 321,306 transcripts per assembly respectively. The BUSCO scores and numbers of orthogroups recovered from orthology analysis of each assembly were both higher on average in the high-quality dataset (Table 1). When we compared the number of transcripts in each individual assembly with the number of orthogroups found for that assembly, we found a significant relationship between these measures in both datasets (linear regression: high-quality dataset, *P* = 0.001; low-quality dataset, *P* = 0.002; Figure 2B). We did not find a significant relationship between the overall *TransRate* scores of assemblies and the number of orthogroups obtained for each assembly (linear regression: high-quality dataset, *P* = 0.43; low-quality dataset, *P* = 0.51; Figure 2D). The number of orthogroups for each dataset was higher in the high-quality dataset, but still largely comparable except for two low-quality read datasets, *Takifugu rubripes* and *Callorhinchus milii*, which each had much lower numbers of orthogroups recovered. In addition to *TransRate* evaluations, the *BUSCO* scores for the low-quality *T. rubripes* and *C. milii* assemblies were also dramatically lower than all other *BUSCO* scores in both datasets (2.7% and 7.2% respectively, compared to the next lowest score: 42.9% for *Notechis scutatus*). However, the overall *BUSCO* scores for the high- and low-quality datasets were not significantly different (Wilcoxon rank sum: *P* = 0.24, Figure 2E). We observed a significant relationship between *BUSCO* score and number of orthogroups recovered in both datasets (linear regression: high quality dataset, *P* = 0.001; low quality dataset, *P* = 0.001; Figure 2F).

**Table 1:** Read set information and transcriptome assembly metrics

| Genus | Accession | Read Length | Number of Reads | High-quality Dataset |  |  |  |  | Low-quality Dataset |  |  |  |  |
| --- | --- | --- | --- | --- | --- | --- | --- | --- | --- | --- | --- | --- | --- |
|  |  |  |  | Number of Transcripts | BUSCO complete | TransRate score | Orthogroups | Species-Specific Orthogroups | Number of Transcripts | BUSCO complete | TransRate score | Orthogroups | Species-Specific Orthogroups |
| *Alligator mississippiensis* | SRR629636 | 100 | 36,130,137 | 287695 | 91.1 | 0.50848 | 12004 | 32 | 466618 | 80.2 | 0.16986 | 10737 | 20 |
| *Ambystoma mexicanum* | SRR5341572 | 101 | 46,417,978 | 209702 | 98.7 | 0.57581 | 11350 | 59 | 528158 | 97.4 | 0.09216 | 10832 | 61 |
| *Anas platyrhynchos* | SRR7127376 | 101 | 20,486,658 | 142201 | 91.1 | 0.65376 | 9813 | 8 | 244848 | 86.8 | 0.2212 | 9129 | 11 |
| *Anolis carolinensis* | SRR391653 | 101 | 17,152,427 | 40327 | 86.2 | 0.33263 | 8729 | 9 | 56207 | 90.1 | 0.18273 | 9093 | 9 |
| *Astyanax mexicanus* | SRR2045431 | 100 | 32,893,691 | 110132 | 98.7 | 0.55641 | 9902 | 44 | 180139 | 97 | 0.21902 | 9187 | 36 |
| *Balaenoptera acutotostrata* | SRR919296 | 100 | 23,923,194 | 200511 | 89.2 | 0.53496 | 11048 | 10 | 364918 | 86.1 | 0.15086 | 9729 | 14 |
| *Bufo bufo* | ERR1331718 | 126 | 37,410,097 | 135770 | 94 | 0.33512 | 11671 | 57 | 413473 | 94.4 | 0.10086 | 11968 | 34 |
| *Caecilia tentaculate* | SRR5591453 | 101 | 28,784,422 | 107413 | 81.8 | 0.56427 | 8737 | 35 | 196546 | 77.9 | 0.15651 | 7993 | 29 |
| *Caiman crocodilus* | ERR2198478 | variable | 31,864,053 | 163595 | 85.8 | 0.28529 | 11113 | 3 | 436573 | 81.5 | 0.20671 | 11475 | 6 |
| *Calidris pugnax* | ERR1018151 | 150 | 13,725,659 | 78074 | 85.5 | 0.46221 | 8239 | 10 | 83535 | 77.9 | 0.24439 | 7880 | 2 |
| *Callorhinchus milii* | SRR513760 | 76 | 35,000,000 | 124415 | 67.3 | 0.32314 | 8418 | 17 | 95463 | 7.2 | 0.15425 | 2232 | 13 |
| *Canis lupus familiaris* | ERR1331673 | 100 | 36,371,999 | 437158 | 83.8 | 0.58601 | 10633 | 3 | 819785 | 86.5 | 0.1697 | 9826 | 10 |
| *Dasyus novemcinctus* | SRR494766 | 101 | 31,705,473 | 55634 | 79.2 | 0.33783 | 8868 | 7 | 192657 | 66 | 0.12049 | 7478 | 6 |
| *Felis catus* | ERR1331679 | 100 | 40,228,662 | 516209 | 79.5 | 0.51854 | 10659 | 8 | 945952 | 85.2 | 0.20215 | 9790 | 4 |
| *Gadhus morhua* | SRR2045420 | 100 | 18,943,673 | 85927 | 74 | 0.46787 | 8082 | 29 | 131171 | 64.3 | 0.21936 | 6919 | 15 |
| *Gallus gallus* | ERR1298598 | 100 | 14,955,711 | 272485 | 72.3 | 0.48137 | 9069 | 8 | 444042 | 62.1 | 0.15068 | 7562 | 9 |
| *Haplochromis burtoni* | SRR387451 | 101 | 16,142,312 | 40240 | 69.3 | 0.34653 | 7981 | 14 | 60824 | 54.5 | 0.19438 | 6379 | 48 |
| *Homo sapiens* | SRR5576267 | 101 | 20,633,201 | 171048 | 72.6 | 0.48352 | 9971 | 5 | 317048 | 74.2 | 0.16465 | 9271 | 8 |
| *Ictalurus punctatus* | SRR917955 | 100 | 28,319,586 | 99232 | 83.8 | 0.49223 | 8645 | 32 | 159608 | 73 | 0.25281 | 7538 | 34 |
| *Latimeria menadoensis* | SRR576100 | 109 | 39,788,120 | 101337 | 73.3 | 0.34696 | 8913 | 69 | 258443 | 82.2 | 0.11311 | 9692 | 13 |
| *Lepidophyma flavimaculatum* | DRR034613 | variable | 20,350,517 | 121895 | 91.4 | 0.28563 | 10505 | 59 | 174935 | 90.7 | 0.14923 | 10395 | 37 |
| *Lepisosteus oculatus* | SRR1287992 | 101 | 22,992,842 | 75239 | 95.4 | 0.44361 | 10235 | 55 | 195782 | 88.5 | 0.15598 | 9172 | 42 |
| *Lethenteron camtschaticum* | SRR3223459 | 125 | 29,559,367 | 125856 | 90.4 | 0.32322 | 8577 | 292 | 274262 | 92.4 | 0.14484 | 8431 | 93 |
| *Lissotriton montandoni* | SRR3299753 | 100 | 32,548,205 | 195142 | 95.4 | 0.46462 | 10934 | 68 | 387445 | 91.1 | 0.12708 | 9939 | 56 |
| *Notamacropus eugenii* | DRR013408,<br>DRR013409,<br>DRR013410 | 100 | 24,378,361 | 198447 | 88.5 | 0.60651 | 9859 | 24 | 347172 | 82.2 | 0.16235 | 8917 | 28 |
| *Notechis scutatus* | SRR519122 | 90 | 25,626,764 | 168738 | 58.7 | 0.43875 | 7908 | 31 | 277137 | 42.9 | 0.18596 | 6254 | 32 |
| *Oophaga sylvatica* | SRR9120851 | 100 | 22,858,029 | 166747 | 56.1 | 0.47685 | 8650 | 18 | 423029 | 49.8 | 0.13789 | 7690 | 24 |
| *Oryctolagus cuniculus* | ERR1331669 | 100 | 22,037,691 | 158880 | 84.8 | 0.5591 | 9304 | 4 | 349879 | 81.8 | 0.11102 | 8469 | 5 |
| *Parus major* | SRR1847228 | 101 | 35,000,000 | 155826 | 95 | 0.54739 | 10349 | 16 | 261539 | 91.7 | 0.20877 | 9408 | 18 |
| *Pelodiscus sinensis* | SRR6157006 | 150 | 24,740,727 | 274343 | 99 | 0.32231 | 12143 | 40 | 367085 | 95.4 | 0.11519 | 11332 | 23 |
| *Pelusios castaneus* | SRR629649 | 100 | 45,163,324 | 254815 | 97.4 | 0.42891 | 12168 | 31 | 419831 | 92.4 | 0.15182 | 10728 | 19 |
| *Protopterus sp.* | ERR2202465 | 150 | 18,298,224 | 327343 | 61.4 | 0.34033 | 9036 | 127 | 141824 | 65.7 | 0.21121 | 8558 | 89 |
| *Rana pipiens* | SRR1185245 | 101 | 35,791,829 | 136439 | 82.2 | 0.52391 | 9695 | 36 | 238110 | 75.4 | 0.20868 | 9029 | 16 |
| *Rhinella marina* | SRR6311453 | 100 | 27,446,915 | 511551 | 67.4 | 0.48377 | 11184 | 48 | 1056698 | 60.4 | 0.16511 | 10330 | 32 |
| *Rhinolophus sinicus* | SRR2273875 | 101 | 30,559,494 | 184384 | 90.8 | 0.68372 | 10658 | 14 | 392613 | 86.2 | 0.1185 | 9933 | 9 |
| *Squalus acanthias* | ERR1525379 | variable | 35,000,000 | 101153 | 84.5 | 0.23542 | 9582 | 25 | 363863 | 83.2 | 0.12189 | 9803 | 79 |
| *Takifugu rubripes* | SRR1005688 | 76 | 35,796,911 | 35375 | 59.7 | 0.44456 | 6518 | 13 | 48271 | 2.7 | 0.1206 | 1287 | 22 |
| *Trachemys scripta* | ERR2198830 | 150 | 22,741,770 | 210713 | 47.8 | 0.48531 | 9322 | 6 | 94129 | 47.6 | 0.16631 | 8123 | 3 |

**Figure 1:**
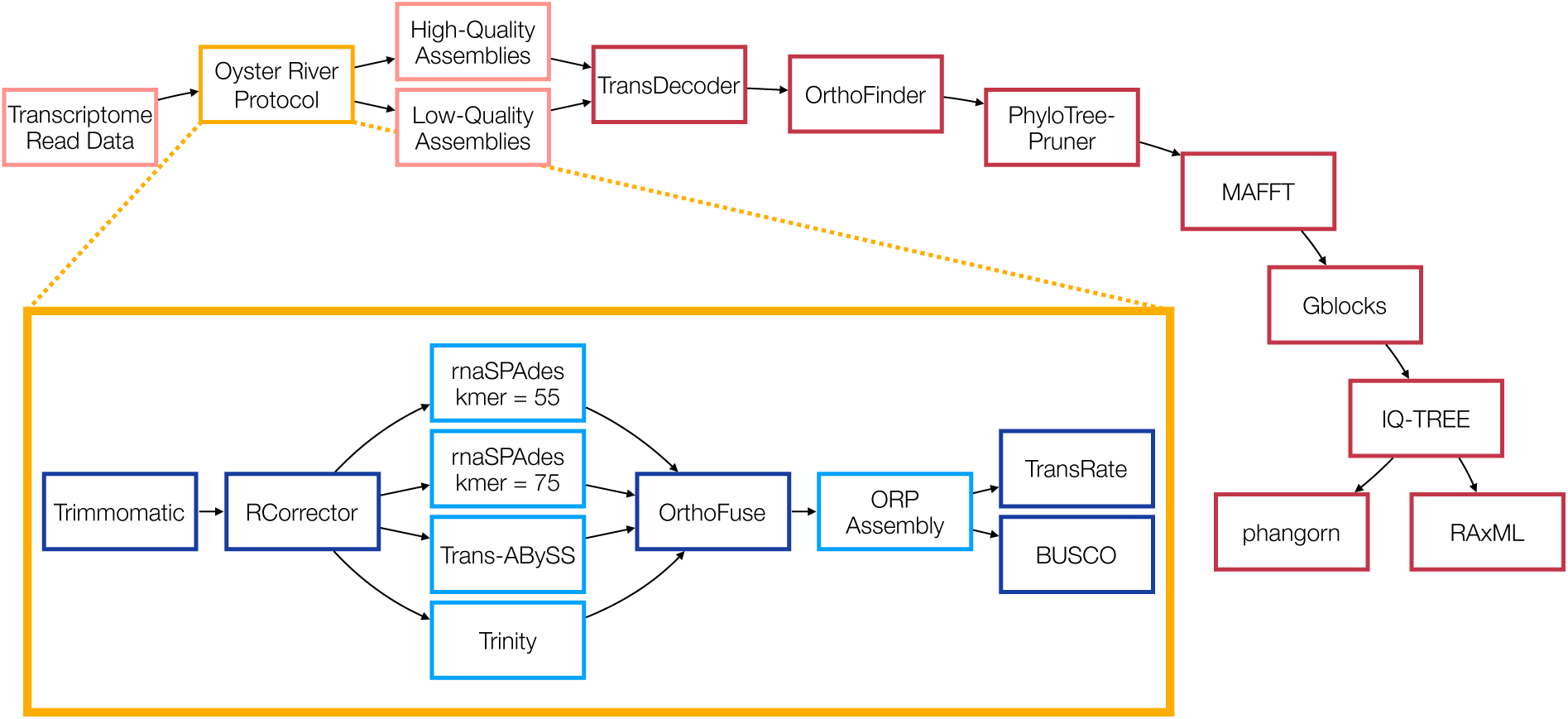
A graphical depiction of the main steps of the phylogenomic pipeline used in this analysis from publicly available transcriptomic datasets to partition tree statistics. In the top flowchart red borders indicate bioinformatic tools used while pink ones depict datasets. The Oyster River Protocol is highlighted in yellow, and in the inset: darker blue borders represent steps of the protocol while the resulting transcriptome assemblies are outlined in lighter blue.

**Figure 2:**
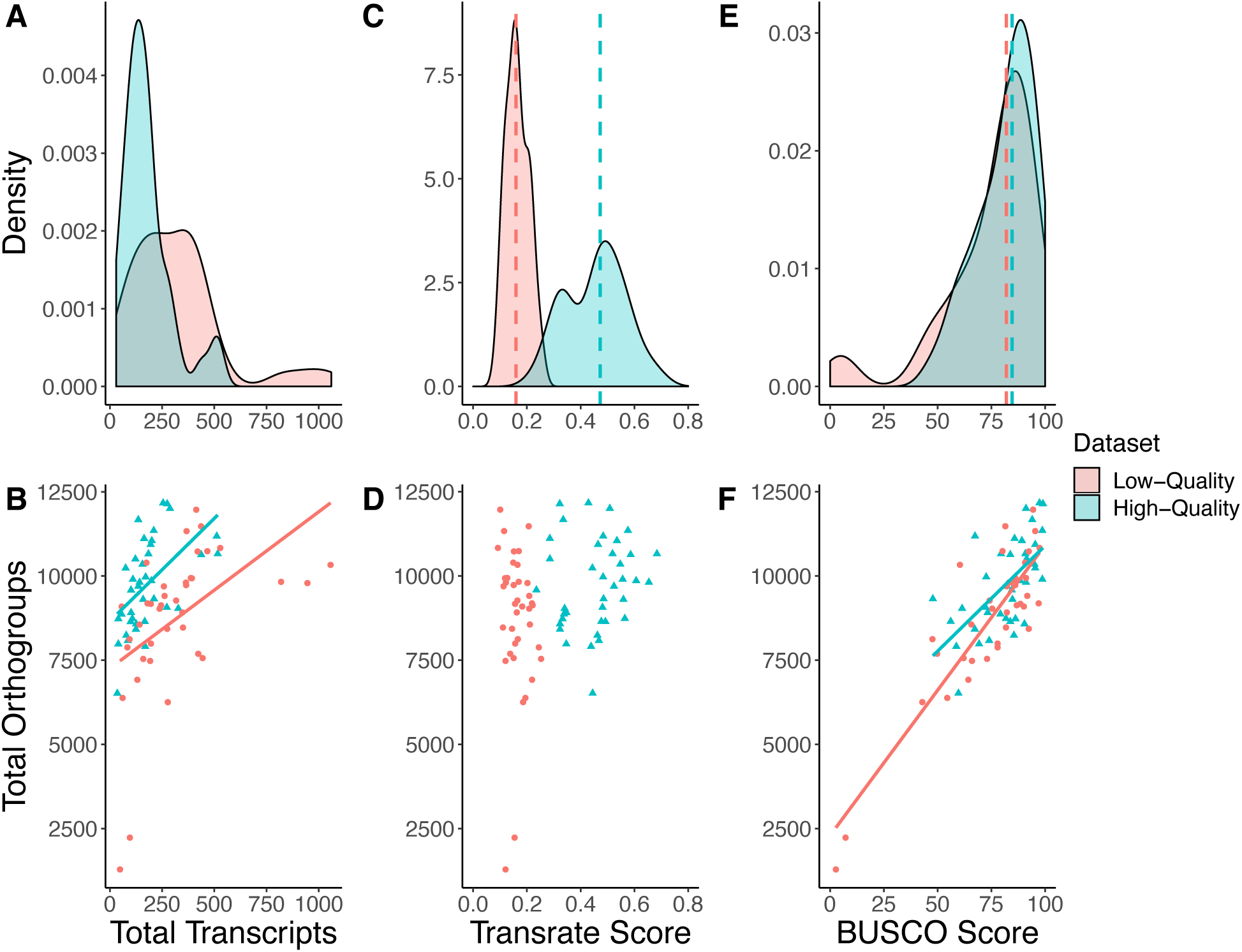
Summary statistics for the high- and low-quality datasets produced. We selected high- and low-quality datasets based on *TransRate* score. This resulted in transcriptome assemblies with both high and low completeness, according to complete *BUSCO* score, in each dataset. Larger assembles in the low-quality dataset did not lead to higher *BUSCO* or *TransRate* scores. Dotted lines in density plots represent medians for each dataset. **A:** Density plot of the total number of transcripts (in thousands) in each transcriptome. **B:** Relationship between the total number of transcripts (in thousands) and the total number of orthogroups. **C:** Density plot of overall *TransRate* scores for each assembly. **D:** Relationship between the overall *TransRate* score and the total number or orthogroups. **E:** Density plot of complete *BUSCO* score for each transcriptome assembly. **F:** Relationship between *BUSCO* score and total number of orthogroups.

### High-quality assemblies result in a larger number of partitions after phylogenomic processing

Next, we isolated one-to-one orthologs that were present in 100% of taxa. After aligning and filtering these orthologs into partitions we observed that one major impact of assembly quality on phylogenomic data matrix construction is the scale of the resulting data. We obtained 2,016 data partitions from the high-quality dataset, whereas we recovered only 408 data partitions from the low-quality dataset. 332 data partitions in both the high- and low-quality datasets included an identical reference sequence from the *Mus musculus* reference transcriptome, demonstrating that a majority of the data partitions recovered from the low-quality dataset are also represented in the high-quality dataset (Figure 3B). The high-quality dataset however, included many more unique sequence partitions (1684 unique partitions compared to 76, Figure 3C). The distributions of alignment lengths between datasets differed significantly before alignment filtering (Wilcoxon rank sum, *P* = 0.02; Figure 3A) with alignments in the high-quality dataset being longer on average, but not after alignment filtering (Wilcoxon rank sum, *P* = 0.79; Figure 3B).

**Figure 3:**
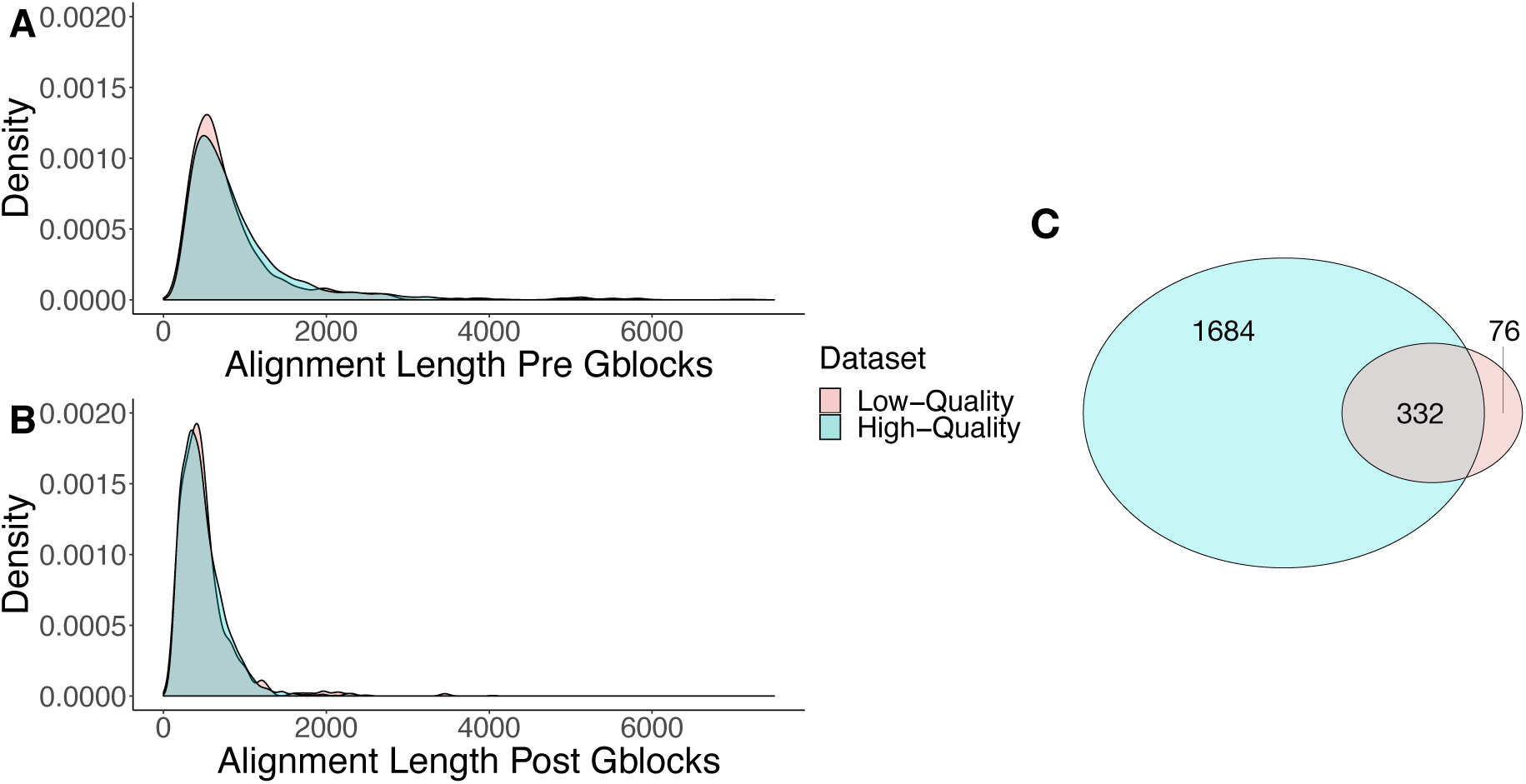
Length of alignments and number of partitions for each dataset. While the lengths of the individual alignments are significantly different before *Gblocks* filtering, they are similar afterwards. The number of partitions recovered through the phylogenomic analysis pipeline is fivefold higher when the dataset is made up of high-quality transcripts compared to lower-quality ones. **A:** Density plot of alignment lengths of each partition before filtering with *Gblocks*. **B:** Density plot of alignment lengths of each partition after filtering with *Gblocks*. **C.** Venn diagram showing number of partitions unique to each dataset, and common between them.

### Less compositional bias and sequence ambiguity in high-quality alignments

In order to draw direct comparisons between the partitions derived from the high- and low-quality datasets, we examined the alignment statistics of the 332 partitions that were shared between them. The percentage of constant sites in each alignment was not significantly different between the high- and low-quality datasets (Wilcoxon rank sum, *P* = .37, Figure 4A). Similarly, the percentage of parsimony-informative sites in the alignments did not differ significantly between the two datasets (Wilcoxon rank sum, *P* = .89, Figure 4B). However, the number of sequences that failed the composition test and the number of sequences with over 50% alignment ambiguity were significantly different between the two datasets (composition – Wilcoxon rank sum, *P* = .006, Figure 4C; ambiguity – Wilcoxon rank sum, P < .001, Figure 4D), and both of these metrics were higher in the low-quality dataset.

**Figure 4:**
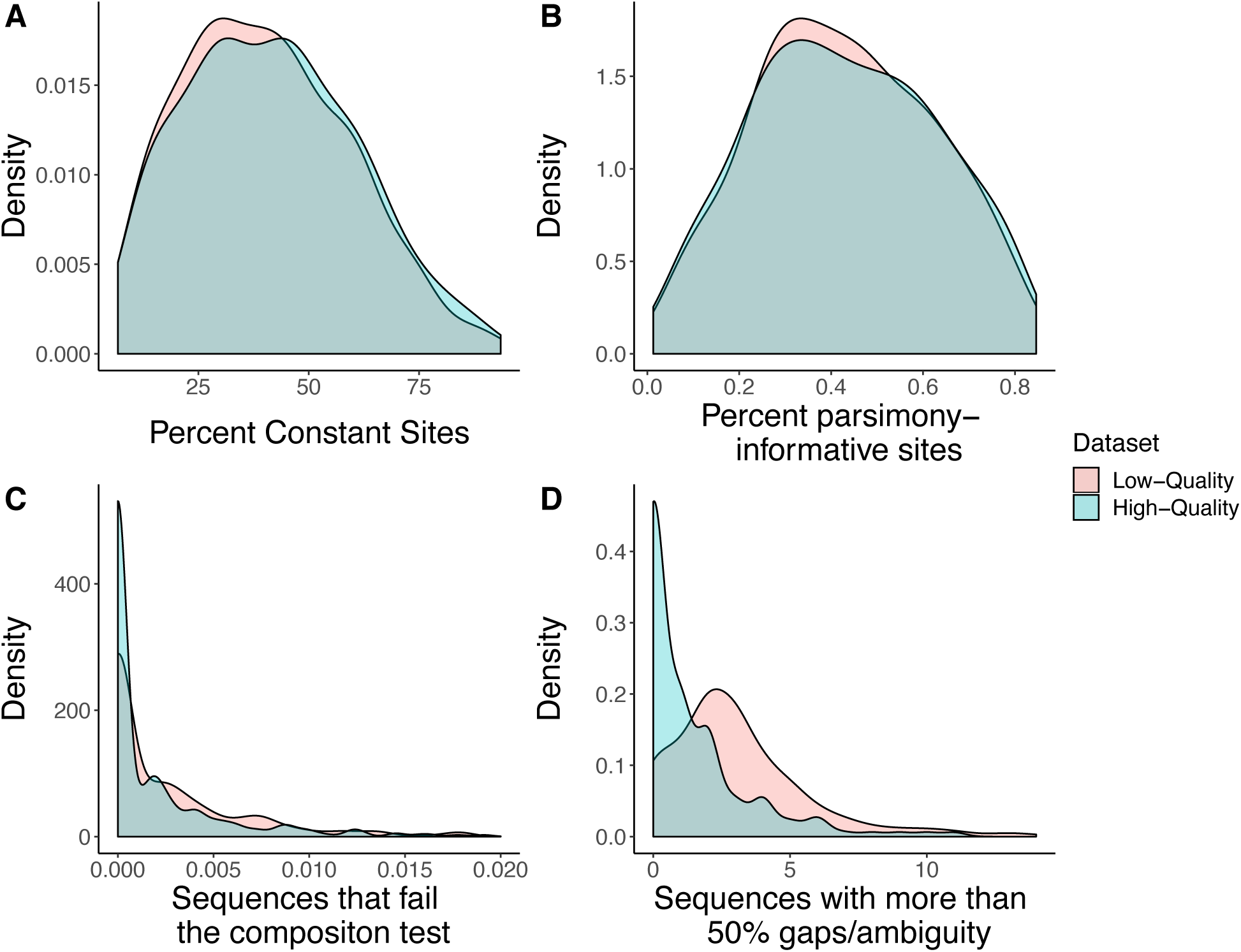
Density plots of four alignment metrics for both datasets. Alignments created from low-quality transcriptome assemblies have similar percentages of constant and parsimony-informative sites, but higher compositional bias and ambiguity when compared to alignments from high-quality assemblies. **A:** Percentage of constant sites in each partition alignment. **B:** Percentage of parsimony-informative sites in each partition alignment. **C:** Number of sequences that fail the composition test, normalized by partition alignment length. **D:** Number of sequences that contain more than 50% gaps/ambiguity in each partition alignment.

### No bias in gene content in partitions from both high- and low-quality datasets

Phylogenetic information content of a given phylogenomic data matrix could be impacted if the partitions themselves are drawn from a biased set of loci. In order to understand the genetic composition of phylogenomic datasets derived from high- and low-quality assemblies, we conducted gene ontology (GO) analysis of the recovered partitions. We did not observe enrichment for functional category in either the high- or low-quality datasets.

### Partitions from high-quality assemblies recapitulate the constraint tree to a larger extent than those from low-quality assemblies

We sought to understand the impact of assembly quality on phylogenetic signal by comparing the two datasets to a constraint tree representing the current view of craniate relationships (Chen et al. 2017; Irisarri et al. 2017) using internode certainty all (ICA) values and Robinson-Foulds (RF) distances. ICA values indicate the proportion of data partitions for the high-quality and low-quality datasets that support each node in our constraint tree (Salichos et al. 2014), whereas RF distances reflect topological differences between partition subtrees and the constraint tree (Robinson and Foulds 1981). The partitions derived from the high-quality dataset possessed characteristically higher ICA values than those from the low-quality dataset, although the distributions of scores were not significantly different (Wilcoxon rank sum, *P* = .47; Figure 5). We found that the high-quality dataset had significantly lower RF values overall than the low-quality dataset (Wilcoxon rank sum, *P* < .001; Figure 6), indicating a shorter distance to the constrained craniate tree for the partitions in the high-quality dataset.

**Figure 5:**
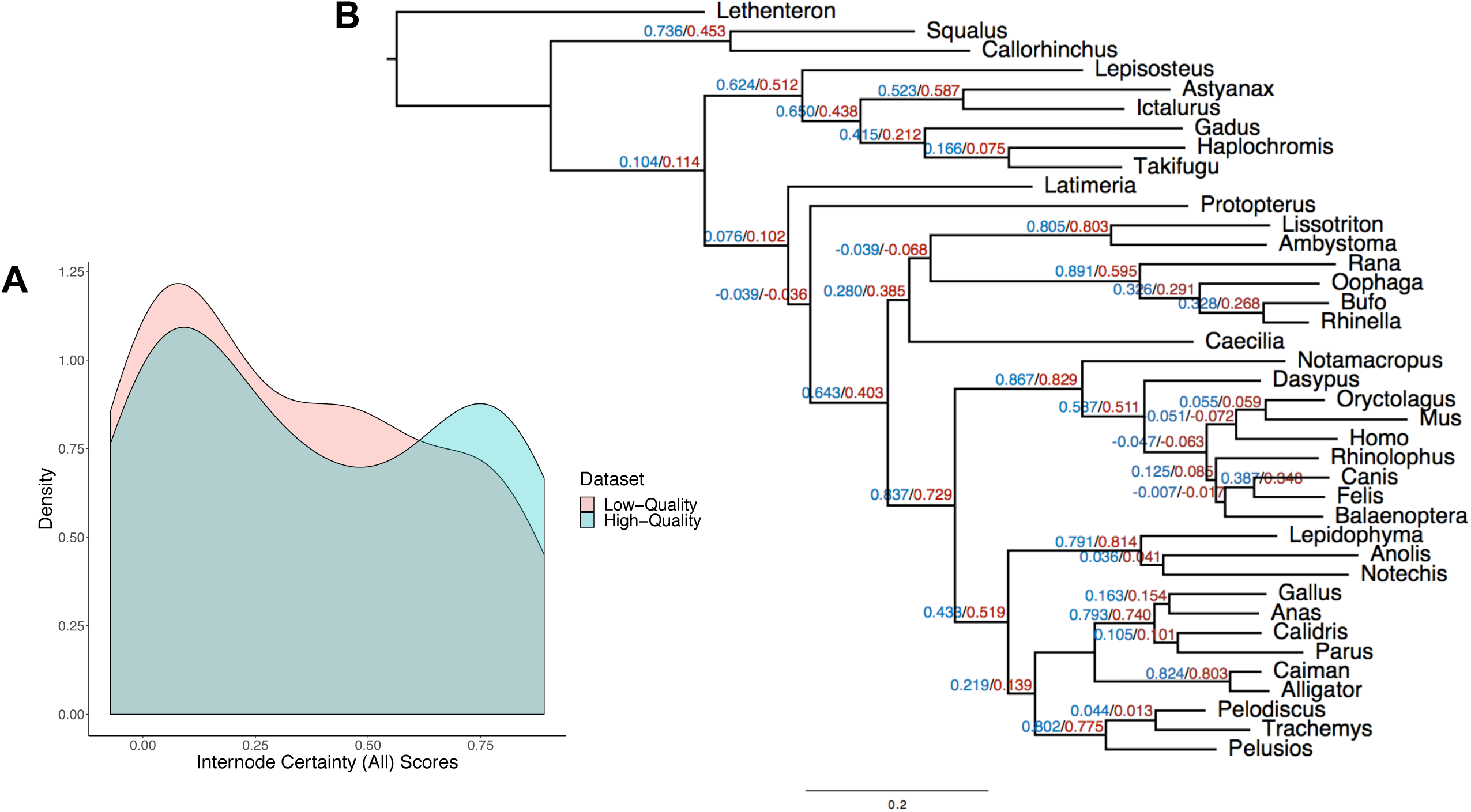
Partitions derived from the high-quality dataset have higher internode certainty all (ICA) values than those derived from the low-quality dataset when compared to the constraint tree. **A:** Density plot of ICA values **B:** Average ICA values for each node. Blue represents the high-quality dataset, red represents the low-quality dataset. Negative ICA values suggest that the node conflicts with at least one other node that has a higher support.

**Figure 6:**
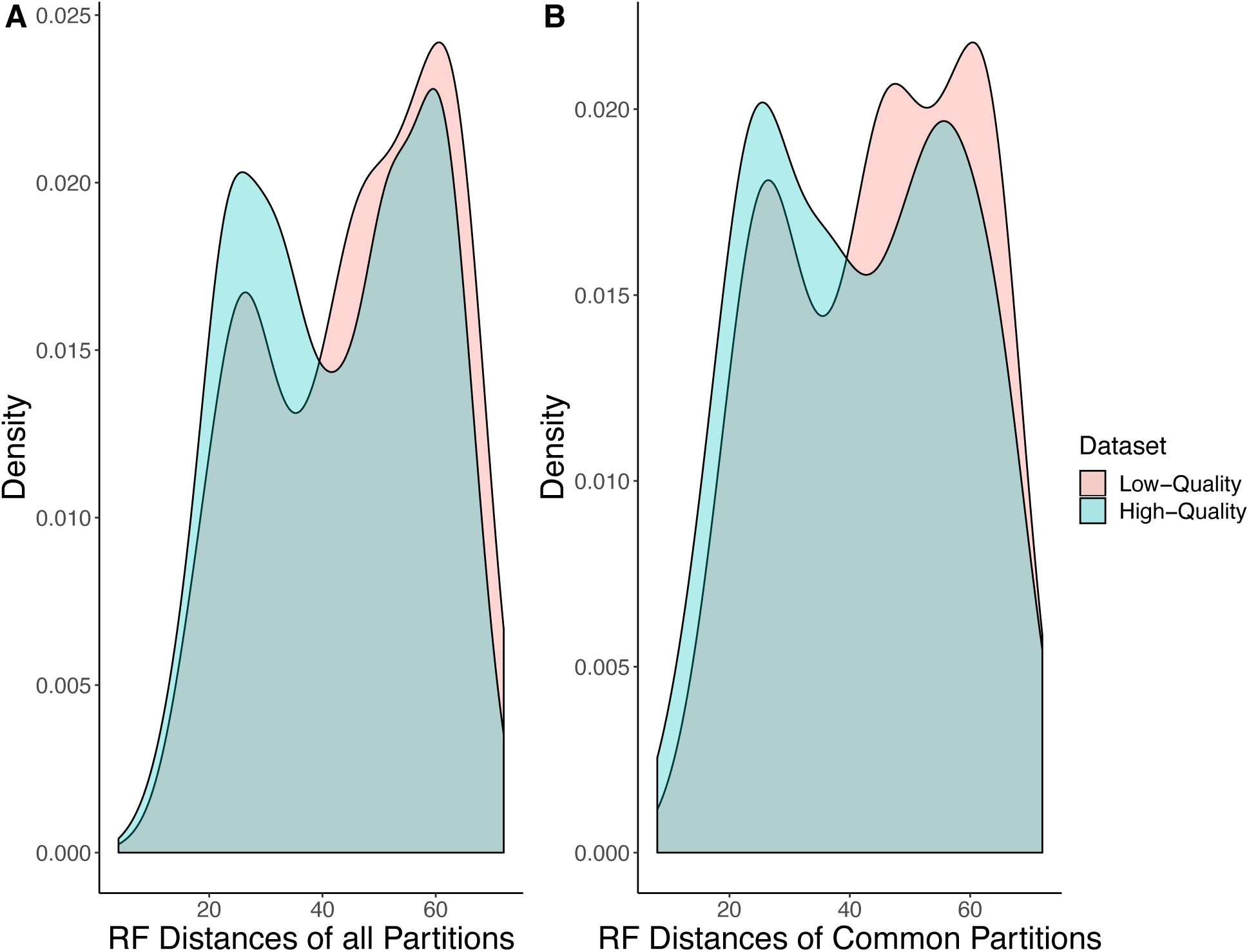
Per partition Robinson-Foulds (RF) distances to the constraint tree are significantly shorter in the high-quality dataset compared with the low-quality dataset. **A:** Density plot for all partitions from both datasets. **B:** Density plot for only those 332 partitions that are shared between the two datasets

In summary, we find that datasets derived from high-quality transcriptome assemblies yield larger phylogenomic matrices than those from low-quality transcriptome assemblies. In addition to being more numerous, the data partitions in the high-quality dataset are also less compositionally biased and have less alignment ambiguity. As a result, the data partitions in the high-quality dataset are more consistent with the craniate phylogeny.

## Discussion

Given the ubiquity of transcriptome usage phylogenomics, we sought to understand how sub-optimal data handing practices during the assembly process may affect downstream phylogenomic analyses. We found that more accurate transcriptome assemblies lead to more accurate phylogenetic reconstructions, likely the product of a richer dataset of orthogroups and a phylogenomic matrix containing more unique data partitions. In addition, high-quality transcriptome assembles contribute to longer alignments, fewer ambiguous regions, less compositional bias, and a greater consistency with the known phylogeny. We did not uncover any functional biases in the GO terms associated with either dataset.

### High-quality assemblies result in a larger number of partitions after phylogenomic processing

The most dramatic difference between the high- and low-quality phylogenomic data matrices is the number of orthogroups that contained all species. After estimating one-to-one orthologs, aligning the orthologs, and filtering the alignment, this difference led to five times the number of data partitions in the high-quality dataset compared with the low-quality dataset.

Transcriptomic assembly errors that are expected to pervade low-quality assemblies include the generation of chimeric transcripts, the generation of incomplete transcripts, or the failure to generate transcripts due to missing data (Smith-Unna et al. 2016; MacManes 2018). Our results from analyses of the low-quality assemblies indicate that incompletely assembled transcripts may be at least partially responsible for the differences in partition number because the partition alignments before filtering are significantly longer in the high-quality dataset, indicating fewer incompletely assembled transcripts in the latter. *OrthoFinder* (Emms and Kelly 2015; Emms and Kelly 2019) may be somewhat robust to these issues, nonetheless when more complete sequence information is provided in high-quality transcripts, *OrthoFinder* analyses identify significantly greater numbers of orthogroups and therefore orthologs. Missing transcripts could also impact the accuracy of downstream analyses and the establishment of one-to-one orthologs because, depending on what data are missing, orthologs and paralogs could become conflated between taxa. Our results are consistent with this expectation because among partitions that are shared between high- and low-quality datasets, those from the high-quality dataset show more accurate phylogenetic signal, as measured by ICA/constraint tree analyses (see below).

We identified two transcriptome assemblies within the low-quality dataset, *Takifugu rubripes* and *Callorhinchus milii*, which have dramatically lower *BUSCO* scores and number of orthogroups recovered than other taxa within the same dataset. We included these two taxa in the analysis despite their extreme *BUSCO* scores for a number of reasons. First, these taxa occupy important phylogenomic positions within the craniate tree and publicly available craniate liver transcriptome datasets are somewhat limited for these branches. Second, while the *TransRate* scores for these two taxa are below average for the low-quality dataset (Figure 2C,D), they are well within the distribution of low-quality assembly *TransRate* scores, indicating that these two taxa have assemblies that are contiguous and correctly assembled to a comparable extent to the other assemblies included in that dataset. While it is standard practice to deposit raw reads into public databases, the read-sets for these two species appeared to have been trimmed prior to public data deposition (Venkatesh et al. 2014), making them shorter than the other read-sets. We identified average read length as the probable reason for the lack of genic completeness as measured by *BUSCO* for these two taxa. Due to this shorter read length, these two organisms performed especially poorly in *rnaSPAdes* with a kmer length of 75 (only reads of length k+1 are used in assembly), which was subsequently the assembly used in the low-quality dataset. Importantly, these two species’ corresponding assemblies in the high-quality dataset were not statistical outliers, indicating that a robust assembly strategy can compensate for sub-optimal sequence reads. Therefore, by including these two taxa, we were able to represent a situation commonly encountered in phylogenomic studies that utilize publicly available data – the inclusion of reads of poor quality or that have been previously processed rather than raw datasets.

The drastic difference in number of partitions in the low-quality dataset compared to the high-quality dataset is due in part to these two taxa having smaller and less complete assemblies than all others. However, when we relax the strict filtering to include orthogroups with up to two missing taxa (thereby giving the low-quality dataset the opportunity to exclude *T. rubripes* and *C. milii*) we find that the high-quality dataset still has over 1600 more partitions than the low-quality dataset, and therefore their inclusion is not the only driving force behind the difference in partitions between the datasets. While there are fewer partitions in the low-quality dataset, it is still a sufficient number for most downstream phylogenomic applications. Therefore, we conclude that while the situation encountered with the *T. rubripes* and *C. milii* RNA-seq data have a disproportionate effect on some aspects of our phylogenomic analysis, their effects are only manifested in analyses of the low-quality assemblies and extend beyond data drop out.

### Low-quality assemblies produce alignments with more compositional bias and alignment ambiguity than high-quality assemblies

In the process of making gene trees for each of the data partitions, *IQ-TREE* calculates a number of metrics about the partition alignments and about the sequences within them (Nguyen et al. 2014). One such test is of compositional homogeneity, which measures the character composition of amino acids in each sequence against the character composition in the whole alignment (http://www.iqtree.org/doc/Frequently-Asked-Questions#what-is-the-purpose-of-composition-test). Heterogeneity or bias in amino acid composition can mislead phylogenetic inferences: distantly-related organisms that have high compositional bias may group together in these analyses (Foster and Hickey 1999). The number of sequences failing the composition chi^2^ test – that is, the number of sequences with higher compositional heterogeneity than expected by chance – was higher in the partitions from the low-quality dataset. Because these partitions have direct counterparts in the high-quality dataset, this difference in compositional heterogeneity is attributable to a difference in assembly quality. Similarly, the partitions from the low-quality dataset also contained more sequences with over 50% gaps or ambiguity in the alignment. While global alignments often contain gaps because of insertions or deletions in the sequences, the comparison of the two datasets implies that these gaps are the result of incorrect transcriptome assemblies rather than natural variation.

The low-quality dataset contained some partitions that the high-quality dataset did not have. These partitions could be unique transcripts only assembled in the low-quality dataset, or they could be the result of differential pruning of paralogous sequences between the two datasets, resulting in a different *Mus* identifying sequence in two partitions that represent the same gene family. They might also be erroneous or duplicate partitions that were misidentified during the *OrthoFinder* procedure as separate gene families due to poorer transcript or assembly quality. In principle, differential data assembly quality could inject bias into the resulting orthogroups if some loci, perhaps short or highly expressed genes, were preferentially assembled among the different datasets, however our GO analyses showed no enrichment or depletion of GO terms in these partitions.

### Partitions derived from high-quality assemblies have higher ICA values and lower RF distances when compared to the constraint tree than those derived from low-quality assemblies

While the ICA values of the high-quality dataset were not significantly higher than those in the low-quality dataset, they did show a greater proportion falling above 0.6. This indicates that the gene trees estimated from the high-quality dataset partitions have more consistency with the constraint tree of craniates than the low-quality dataset. We also showed that the high-quality dataset was closer to the topology of the constraint tree by calculating RF distances. The high-quality dataset had significantly smaller RF distances to the constraint tree (Wilcoxon rank sum, *P* < .001), indicating that the partition trees from the high-quality dataset were more concordant with the constraint tree than those from the low-quality dataset. Since phylogenetic signal for a gene family is a measure of how closely that gene tree matches the species tree (Revell et al. 2008), we also conclude that the high-quality dataset partitions have greater phylogenetic signal than their low-quality counterparts, even when only partitions represented in both datasets are taken into account (Figure 6B). Therefore, our results indicate that a smaller number of well-assembled data partitions have more signal to resolve phylogenies than an equal or greater number of poorer-quality partitions.

### Conclusions

Phylogenomic approaches leverage great power to resolve phylogenetic relationships, but they also include many analytical pitfalls associated with ortholog identification, alignment filtering, and model selection. While these pitfalls have been well-characterized, we chose to focus on transcriptome assembly quality – a more fundamental and largely overlooked aspect of phylogenomic analyses. We addressed this problem empirically using a study design that controls for variables including taxon selection, data type, data provenance, and phylogenetic uncertainty. We show that assembly quality, when all other factors are controlled, can have a dramatic impact on phylogenomic analyses in three ways. First, the richness and size of the dataset can differ profoundly when assembly errors are prevalent in the data. Second, alignments created from low-quality assemblies are more prone to ambiguity and compositional bias than their high-quality counterparts. And third, the partitions derived from high-quality assemblies have greater phylogenetic signal and a significantly greater power to resolve true evolutionary relationships than partitions derived from low-quality assemblies. We conclude that additional analytical interventions aimed at improving assembly quality, such as the Oyster River Protocol (MacManes 2018), are likely worth the additional effort.

## Methods

### Read selection and assembly

To understand the effects of transcriptome assembly quality on phylogenomic inference, we created two datasets, one of high and one of low quality, from publicly available transcriptomic reads. All read data are available on the European Nucleotide Archive (Table 1). We focused on craniates because there are few remaining disputes on the craniate phylogeny (Irisarri et al. 2017) and these well-established phylogenetic relationships serve as a comparison to the topologies found using our high- and low-quality transcriptome assemblies. Our research strategy was to assemble high- and low-quality transcriptomes from the same set of reads. We obtained Illumina-generated paired-end liver transcriptomic reads for 37 vertebrate species spanning the majority of the diversity contained within the clade as well as one craniate outgroup. We assembled each read set using the Oyster River Protocol (ORP) version 2.2.3 (MacManes 2018) on a Linux computer with 24 CPUs and 128GB of RAM. In brief, this protocol begins by adapter- and quality-trimming reads using *Trimmomatic* version 0.38 (Bolger et al. 2014) as per recommendations in MacManes (2014), after which it corrects read errors using *Rcorrector* version 1.0.8 (Song and Florea 2015) following recommendations from MacManes and Eisen (2013). The ORP then assembles trimmed and corrected reads using three different assemblers: *Trinity* version 2.8.5 (Haas et al. 2014) with a kmer length of 25, *Trans-ABySS* version 2.0.1 (Robertson et al. 2010) with a kmer length of 32, and *rnaSPAdes* version 3.14 (Bushmanova et al. 2019) using kmer lengths of 55 and 75. The protocol continues by merging the resultant four assemblies and clustering them into isoform groups. The ORP then scores all transcripts using *TransRate* version 1.0.3 (Smith-Unna et al. 2016) which maps the read sets onto the assembly and assigns quality scores based on the mapping to each transcript and to the assembly as a whole. The protocol selects the member of each group with the highest *TransRate* score and places it into a new file. Finally, the ORP uses *cd-hit-est* version 4.8.1 (Li and Godzik 2006) and a 98% sequence identity threshold to reduce transcript redundancy. The assemblies produced by the ORP are therefore populated by the highest quality, non-redundant sequences produced by any of the five possible assembly strategies (MacManes 2018). A graphical summary of this protocol and our phylogenomic pipeline can be found in Figure 1.

### Quality analysis and high- and low-quality dataset construction

We evaluated each of the five assemblies generated from the ORP (from *Trinity, TransABySS, rnaSPAdes* at two kmer lengths, and the final ORP assembly) for each species in two main ways. We used *BUSCO* version 3.0.1 (Simao et al. 2015), which uses benchmarking universal single copy orthologs to measure the genic completeness of an assembly. In addition, because we were primarily interested in assessing the structural differences in the transcriptome assemblies arising from errors during the assembly process, we generated *TransRate* scores for each assembly, choosing the assembly with the highest overall *TransRate* score for each species to be part of the high-quality dataset, and the one with the lowest overall score to be part of the low-quality dataset. We performed all subsequent steps on both datasets in parallel.

### Orthogroup inference, statistics, and data partition creation

We used *TransDecoder* version 5.5.0 (Haas & Papanicolaou; https://github.com/jls943/TransDecoder) to translate all transcript sequences to amino acid sequences. The transcriptome assembly process assigns each new transcript a unique name so that it can be differentiated within the assembly. This means that the high- and low-quality assemblies do not share identical transcripts or names common to both assemblies, making the direct comparison of sequences impossible. To circumvent this issue, we added the *Mus musculus* reference transcriptome (release 96) (Howe et al. 2020) to both datasets just before the *TransDecoder* step so that a *Mus* sequence would be present in many orthogroups and partitions downstream. This created a common naming system by which we could compare the content of orthogroups and partitions derived from assemblies of high and low quality later in the analysis.

For each dataset (containing either the high-quality or low-quality transcriptome assemblies for the 38 craniate species plus the *Mus* reference transcriptome) we performed a separate *OrthoFinder* version 2.3.3 analysis (Emms and Kelly 2015; Emms and Kelly 2019). We then used linear regressions in *R* version 3.5.2 (R Core Team 2018) to evaluate the relationship between the total number of orthogroups found for each taxon and three other measures: the total number of transcripts in each assembly, the overall *TransRate* score, and the *BUSCO* complete score. We also plotted the distributions of these three measures for each dataset and performed Wilcoxon rank sum tests in *R* to determine if they were statistically different.

We filtered the resulting orthogroups so that we retained only those that had each taxon represented by at least one sequence. From these, we obtained one-to-one orthologs using *PhyloTreePruner* (Kocot et al. 2013). We realigned these sequences using *MAFFT* version 7.305b using the “auto” setting (Katoh and Toh 2010), and filtered the alignments for poorly aligned or divergent regions with *Gblocks* version 0.91b (Castresana 2000; Talavera and Castresana 2007). Finally, we concatenated all sequences into a NEXUS file for each dataset. We measured the lengths of the alignments both before and after *Gblocks* and compared the content of both groups of partitions by using the *Mus* sequence headers as common identifiers that were present in both datasets and determined the numbers of unique and shared partitions. We then used *IQ-TREE* version 1.6.12 under the LG model (Nguyen et al. 2014) to find individual gene trees for each partition in each dataset.

### GO analysis and alignment metrics

To investigate the differences in content and qualities of the partitions between the two datasets, we separated the partitions into groups containing only those that were unique to each dataset, and only those that were shared between the two datasets. We used *InterProScan* version 5.31-70.0 (Jones et al. 2014) to annotate the partitions unique to each dataset and then performed a gene ontology (GO) analysis with *topGO* version 2.32.0 (Alexa and Rahnenfuhrer 2009) in *R* version 3.5.2 (R Core Team 2018) to check for any functional enrichment or depletion bias in the partitions of either dataset. For each partition common to both datasets, we extracted various alignment metrics from the log and information files generated while making partition trees in *IQ-TREE*. These included percent constant sites, percent parsimony-informative sites, number of sequences that failed the chi^2^ composition test (which we normalized by alignment length), and the number of sequences that contained more than 50% gaps or ambiguity. To test for significant differences, we performed Wilcoxon rank sum tests in *R* version 3.5.2 (R Core Team 2018) between the two datasets for each of these measures.

### Constraint tree and comparisons of partition trees

The phylogenetic relationships among the 38 craniate species for which we obtained liver RNA-seq data are well-supported by previous work (Irisarri et al. 2017). Therefore, we used a tree that reflects the most well-supported hypothesized relationships for comparison against the partition trees. Using *Mesquite* version 3.6 (Maddison and Maddison 2018), we constructed a constraint tree that reflects the widely accepted topology for craniates. We used the high-quality dataset NEXUS alignment file along with this topology to estimate the constraint tree topology with branch lengths in *IQ-TREE* using the LG model (Nguyen et al. 2014). We used *RAxML* version 8.2.11 (Stamatakis 2014) to calculate ICA values between the individual partition trees and the constraint tree. The ICA refers to the degree of certainty for each internal node of the tree compared to the constraint tree when all other conflicting bipartitions are taken into account for that dataset. Numbers close to 1 show a lack of conflict between the partition tree and the constraint tree (Salichos et al. 2014). We also calculated RF distances (Robinson and Foulds 1981) from the partition trees in each dataset to the constraint tree using *phangorn* version 2.5.5 (Schliep 2011) in *R* version 3.5.2 (R Core Team 2018). This metric likewise measures the differences in topology (RF distance) from the partition trees to the constraint tree, but in this case, smaller numbers indicate less conflict between the two trees. We tested for significant differences between the two dataset distributions using a Wilcoxon rank sum test in *R* version 3.5.2 (R Core Team 2018) for both ICA values and RF distances.

### Data availability and Reproducibility

Assemblies used in this study are available here: https://doi.org/10.5281/zenodo.3939160. All custom scripts written for or used in this work as well as commands for programs run are accessible via the GitHub repository: http://github.com/jls943/quality_review.

## Acknowledgements and Funding

JLS, DCP, and MDM were supported through NSF grant No. 1638296. MDM was additionally supported by NIH award No. 1R35GM128843. DCP was additionally supported by NSF grant No. 1755337. We thank Joseph Ryan and Sabrina Pankey for their helpful comments, and Toni Westbrook for his unending assistance.

